# ELDR-Glo, a biosensor for cell age that correlates with quiescence depth

**DOI:** 10.64898/2026.07.04.736063

**Authors:** Martha S. Johnson, Sapna Kamath, Dalia Fleifel, Tessa Hill, Liu Mei, Neha Das, Mary Linares, Wenyih Aw, Victoria L. Bautch, Jeanette Gowen Cook

**Affiliations:** Department of Biochemistry and Biophysics; Department of Biochemistry and Molecular Biology, University of Nebraska Medical Center; Department of Biology, University of North Carolina, Chapel Hill, NC 27599-7260; Joint Department of Biomedical Engineering, University of North Carolina, Chapel Hill, NC 27599-7260; Department of Pharmacology, University of North Carolina, Chapel Hill, NC 27599-7260; Lineberger Comprehensive Cancer Center, University of North Carolina, Chapel Hill, NC 27599-7260

## Abstract

Fluorescent reporters are powerful tools for revealing intercellular heterogeneity among proliferating cells. However, there are few tools to analyze differences among quiescent (G0) cells, though such differences are relevant for development, tissue maintenance, and cancer cell behavior. Quiescence heterogeneity, also known as quiescence depth, typically correlates with time after cell cycle arrest, yet directly measuring cell age is not feasible for all cell types or most tissues. Here, we describe ELDR-Glo, a genetically-encoded fluorescent biosensor that estimates relative cell age, or time since the last cell cycle. The biosensor integrates replication-coupled degradation in S phase with a slow-maturing mCherry and a normalization module. We demonstrate that ELDR-Glo signal correlates with true cell age by both live-cell imaging and in fixed cells. ELDR-Glo distinguishes early and late G0 cells and functions as a relative quiescence depth reporter *in situ*. The biosensor is compatible with multiplexed immunofluorescence and flow cytometry. ELDR-Glo provides a unique and scalable tool to investigate cell proliferation control.

**Teaser:** A fluorescent biosensor to distinguish cells of different ages and thus, likelihood of returning to proliferation from quiescence.

## Introduction

Precise regulation of cell proliferation is essential for development, tissue homeostasis, and regeneration, whereas dysregulation is a hallmark of tumorigenesis and cancer (reviewed in (*1–5*)). Proliferation requires progression through cell cycle phases known as G1 (gap 1), S (DNA synthesis), G2 (gap 2), and M (mitosis). Cells can exit the cell cycle in response to various external and internal cues into a reversible, non-dividing state known as quiescence or G0. In this state, cells have altered metabolic and transcriptional properties, yet they retain the capacity to re-enter the cell cycle after appropriate stimulation (reviewed in (*6–10*)).

Quiescence is not a uniform state, however. Cells in shallow quiescence require less stimulation and less time to re-enter the cell cycle, whereas cells in deeper quiescence require stronger or more prolonged stimulation to resume proliferation (*11, 12*). Molecular differences between deep and shallow quiescent cells include tissue-specific factors in tissue stem cells and lysosome function or chromatin modifications in fibroblasts (*13–15*). Currently, measuring quiescence depth involves re-stimulating a population of arrested cells and monitoring the timing and frequency of cell cycle re-entry. As a result, one can infer after the fact how deeply quiescent a cell *had been* before it was stimulated, but not how deeply quiescent a cell *is* before stimulation.

There is a strong correlation between how long cells have been arrested and how deeply quiescent they are. In other words, older cells - those have spent more time since their last cell cycle - are generally more deeply quiescent than younger cells (*16, 17*). Unlike markers of tissue or organismal age such as senescent cell biomarkers (*18*), there are few available molecular markers of individual cell age. One very specific example is the decrease in acetocholinesterase as red blood cells age (*19*), but there is no apparent general marker or reporter of cell age.

Many methods can identify cell cycle phase by molecular markers and cellular DNA content. A variety of tracing methods can also report the number of cell divisions over time both *in vitro* and *in vivo* (*20*). Live cell imaging with genetically-encoded fluorescent biosensors has become a powerful tool to capture precise dynamics during cell cycle progression and the status of key cell cycle activities (*21–26*). Cell age can be directly measured by long-term live cell imaging, but is restricted to appropriate cultured cell types, specific organisms, and accessible tissues (*27–29*). There are studies relevant to both normal and abnormal biology that could benefit from a method to distinguish older and younger cells without the need for live cell imaging or cell type-specific markers.

Knowing relative cell age would provide critical insight into cell fate decisions, including the likelihood of cell-cycle re-entry (*17*) or differentiation (*30*), and this information is highly relevant to understanding development, aging, and cancer cell behavior. It would also enable more precise interpretation of heterogeneous populations, where cells occupying the same state may differ markedly in their functional potential.

To address this need, we developed a genetically-encoded fluorescent biosensor based on a slow-maturing mCherry (*31*). This biosensor converts absolute time since the last cell cycle into a quantitative fluorescence signal. We characterized ELDR-Glo dynamics in multiple settings as a tool to estimate cell age and to distinguish recently-arrested cells from cells that have been arrested for much longer. We show that the biosensor signal correlates with quiescence depth. ELDR-Glo is also compatible with multiplexed imaging and flow cytometry. By enabling scalable, snapshot-based measurements of temporal cell state, ELDR-Glo addresses a unique gap in cell cycle technology.

## Results

### ELDR-Glo design and expression

Our goal for the biosensor is a fluorescence intensity signal that correlates with cell age. Toward that end, we constructed a genetically-encoded timer using an mCherry variant that has slow fluorescence maturation kinetics first described by Subach *et al*., (*31*). We fused this variant to a replication-coupled destruction motif to induce profound degradation during each S phase (Figure 1A). The mCherry protein should begin to re-accumulate in G2 but be most abundant in G1 phase and G0 (Figure 1B). Because it accumulates in G1 and G0, we named the biosensor **E**mits **L**ight in **D**ormancy as a **R**eporter of **G**1/G0 (ELDR-Glo).

**Figure 1.**
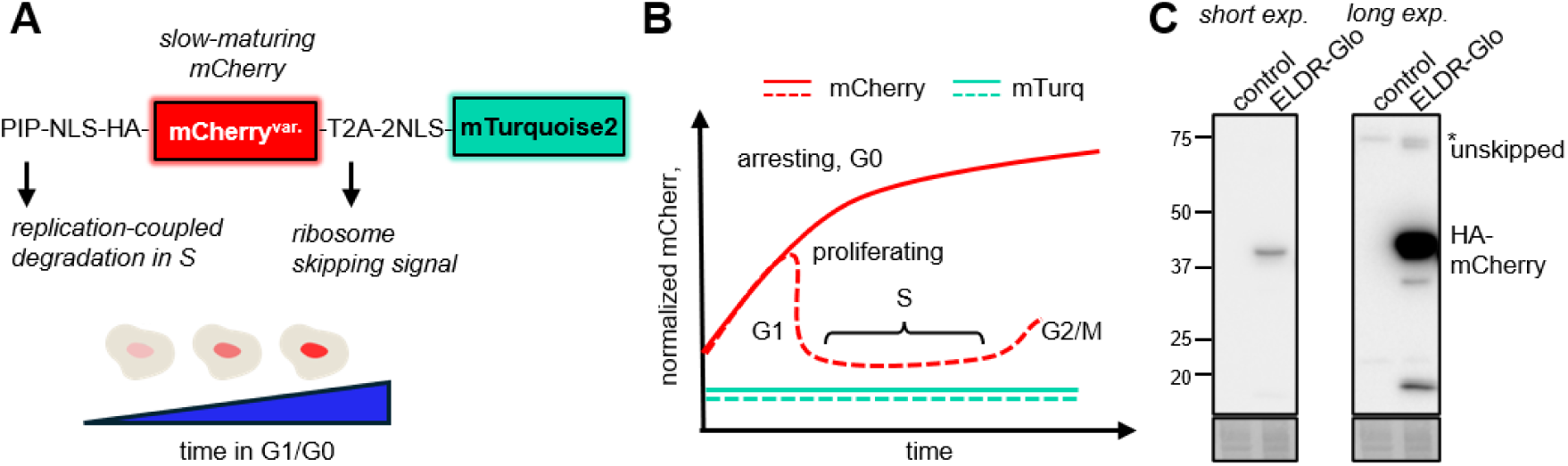
Design and expression of the ELDR-Glo cell age reporter. (A) Schematic of the ELDR-Glo construct. The reporter contains the PIP degron from human CDT1 (aa 1–17) fused to an HA-epitope, nuclear-localization signal and mCherry fluorescent timer followed by a T2A ribosomal skipping sequence and two nuclear localization signals fused to mTurquoise2. (B) Model of ELDR-Glo dynamics across the cell cycle. Solid lines are arresting cells, and dashed lines are proliferating cells. (C) Anti-HA immunoblot of ELDR-Glo expression in parent and ELDR-Glo RPE cells. The position of presumed unskipped fusion is marked; asterisk is a background band. Short and long exposure images are shown; total protein by Ponceau S staining as loading controls.

The ELDR-Glo biosensor contains the PCNA interacting peptide (PIP) degron from the human CDT1 origin licensing protein (amino acids 1–17). This PIP degron is sufficient to target proteins for replication-coupled destruction during S phase via the CRL4^CDT2^ E3 ubiquitin ligase (*32*). We had previously characterized a similar fusion to fast-folding fluorescent proteins and quantified their rapid degradation at the onset of S phase (*21*); a related fusion has also been well-described (*23.*) We fused this degron, a nuclear localization signal, and an HA epitope tag to a slow-maturing mCherry variant. We stably expressed the fusion from a recombinant lentiviral vector in a human non-transformed epithelial cell line, RPE1_hTert (hereafter “RPE”).

Although the mCherry variant undergoes intrinsic blue-to-red fluorescence conversion during maturation (*31*), we did not detect the blue intermediate form at this expression level under live-cell imaging conditions. Therefore, to normalize for intercellular variations in expression, we co-expressed a fast-folding, stable, nuclear mTurquoise2 via a T2A ribosome skipping sequence (Figure 1A and Methods). We confirmed efficient T2A cleavage and biosensor expression by immunoblotting for the HA epitope tag. We detected a primary band corresponding to the expected PIP–NLS-HA–mCherry product, with almost no detectable unskipped fusion protein (Figure 1C). (Any unskipped fusion also contains the PIP degron for S phase degradation and thus would not confound interpretation of the mCherry/mTurquoise signal.)

### ELDR-Glo dynamics in proliferating cells

To visualize ELDR-Glo dynamics, we performed long-term live-cell fluorescence imaging in RPE cells co-expressing a proliferating cell nuclear antigen (PCNA) fusion as a marker of S phase (*26*) (see Methods). During S phase, PCNA concentrates into characteristic nuclear foci that correspond to sites of DNA synthesis, so we used PCNA-mVenus foci formation and dissolution to identify entry into and exit from S phase. We used semi-automated tracking of cells expressing ELDR-Glo and quantified the ratio of mCherry fluorescence to mTurquoise fluorescence (“normalized mCherry”) relative to time and relative to cell cycle transitions. PCNA foci formation coincided with rapid loss of ELDR-Glo mCherry signal, but not mTurquoise, as expected; a representative cell is shown in Figure 2A and the full trace of one mother-daughter pair in Figure 2B. G1 phase was slightly longer in this daughter than in the mother, so ELDR-Glo signal at the end of G1 was slightly higher in the daughter (Figure 2B). After S phase, normalized mCherry signal began to increase again. (The signal increase around cell division is the pair of G1 daughters.) After tracking many live cells, we used linear regression to calculate the average rate of increase during G1 phase (Figure 2C). We also pulse-labeled cells with the thymidine analog, EdU, then imaged after fixing proliferating or arrested cells as a control. ELDR-Glo signal was low in EdU-positive S phase cells, but of varying intensities in EdU-negative G1 and G2 cells (Supplementary Figure S1). We observed the same increase during G1 phase in multiple other cell lines, including the breast cancer cell line, MCF-7 (Supplementary Figure S2)

**Figure 2.**
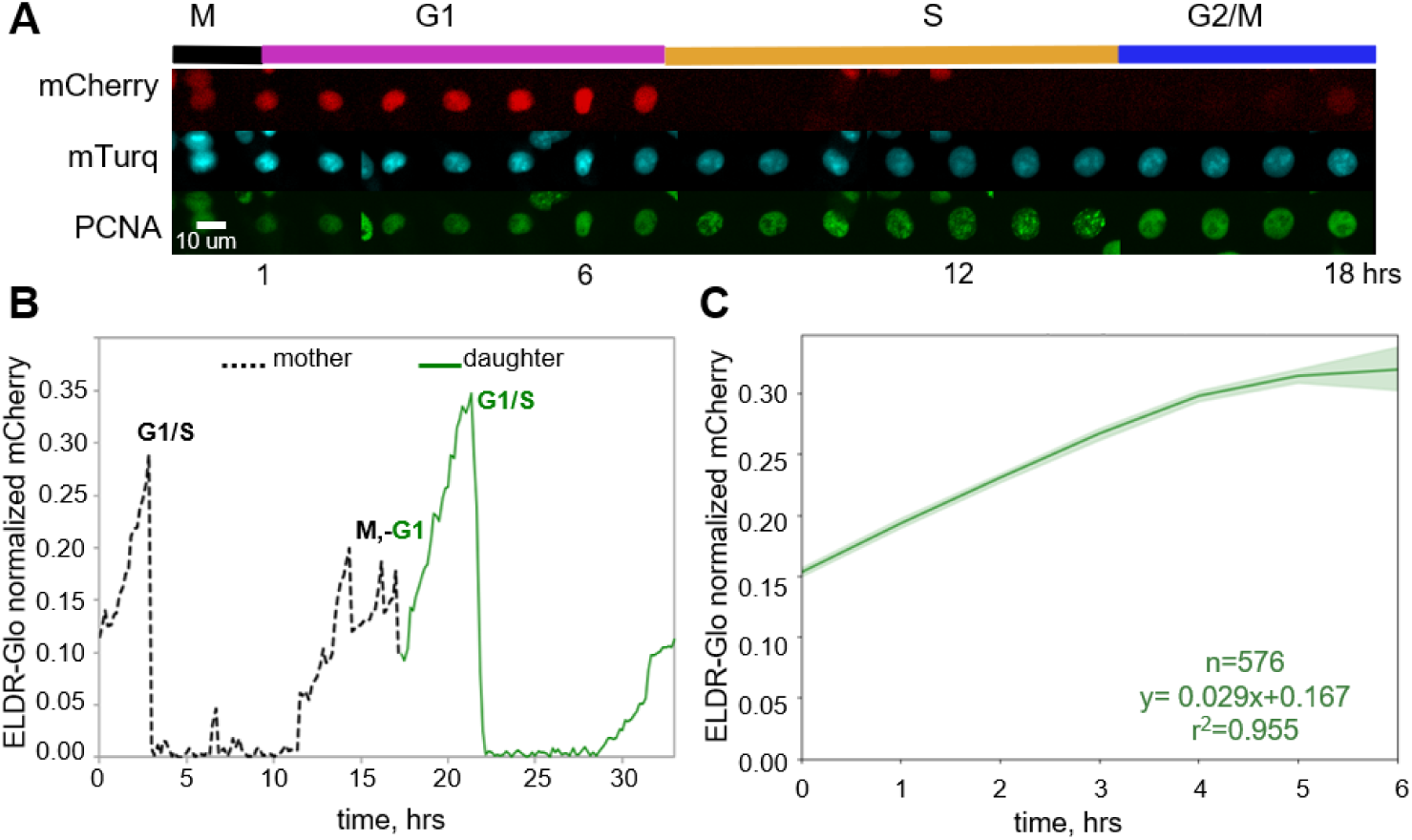
ELDR-Glo dynamics by live cell imaging of proliferating cells. (A) Representative time-lapse filmstrip one frame per hour of a proliferating RPE cell expressing ELDR-Glo and PCNA-mVenus. (B) Single-cell mother–daughter fluorescence trace showing normalized mCherry intensity (mCherry/mTurquoise) for cells expressing ELDR-Glo. “M” indicates the cell division event into two G1 daughter cells. G1/S indicates the transition from G1into S phase. (C) Median normalized mCherry traces for ELDR-Glo in G1 phase from mitosis to S phase onset in 576 tracked cells. Shaded regions represent the 95% confidence interval.

PIP degrons can also mediate protein degradation after substantial DNA damage (*33, 34*). We tested if endogenous DNA damage or replication stress contribute to biosensor dynamics, which would confound interpretations of ELDR-Glo signal. We treated cells with low to moderate concentrations of the DNA damaging agent neocarzinostatin (NCS). At concentrations that were sufficient to induce a robust DNA damage response evidenced as 53BP1 foci, ELDR-Glo signal was not substantially affected (Supplementary Figure S3). In contrast, S phase cells reduced ELDR-Glo signal to a much larger degree than even the highest tested NCS dose, i.e., S phase degradation was much stronger than DNA damage (Supplementary Figure S3C). We are thus confident that ELDR-Glo dynamics are attributable to cell cycle progression and not spontaneous endogenous DNA damage.

### ELDR-Glo dynamics in arresting cells

We next examined ELDR-Glo dynamics during cell cycle exit. We induced arrest with palbociclib, a pharmacological inhibitor of the G1 kinases, CDK4 and CDK6. We treated cells at the beginning of live cell imaging and tracked them from their final division. After division in the presence of the drug, daughter cells arrested and accumulated ELDR-Glo signal continuously (example in Figure 3A). Because these cells did not enter S phase, their signals increased to much higher levels than the maximum G1 signal in typical proliferating cells (compare to Figure 2). In arresting cells, normalized mCherry intensity increased linearly before approaching a plateau beyond which there was less increase with time. This slow-down occurred approximately 40-50 hours after cell division in these cells under these conditions (Figure 3B). We observed similar increases in cells arrested by mitogen withdrawal, contact inhibition, or CDK4/6 inhibitor treatment (Supplementary Figure S4A-C). Interestingly, the rate of signal increase was somewhat slower in mitogen-starved cells (Figures S4C) which may reflect reduced protein synthesis during this type of arrest(*35*). We also observed linear signal increases during arrest in CDK4/6 inhibitor-treated MCF-7 cells and in human umbilical vein endothelial cells (HUVEC) arrested by contact inhibition (Supplementary Figure S4D-F).

**Figure 3.**
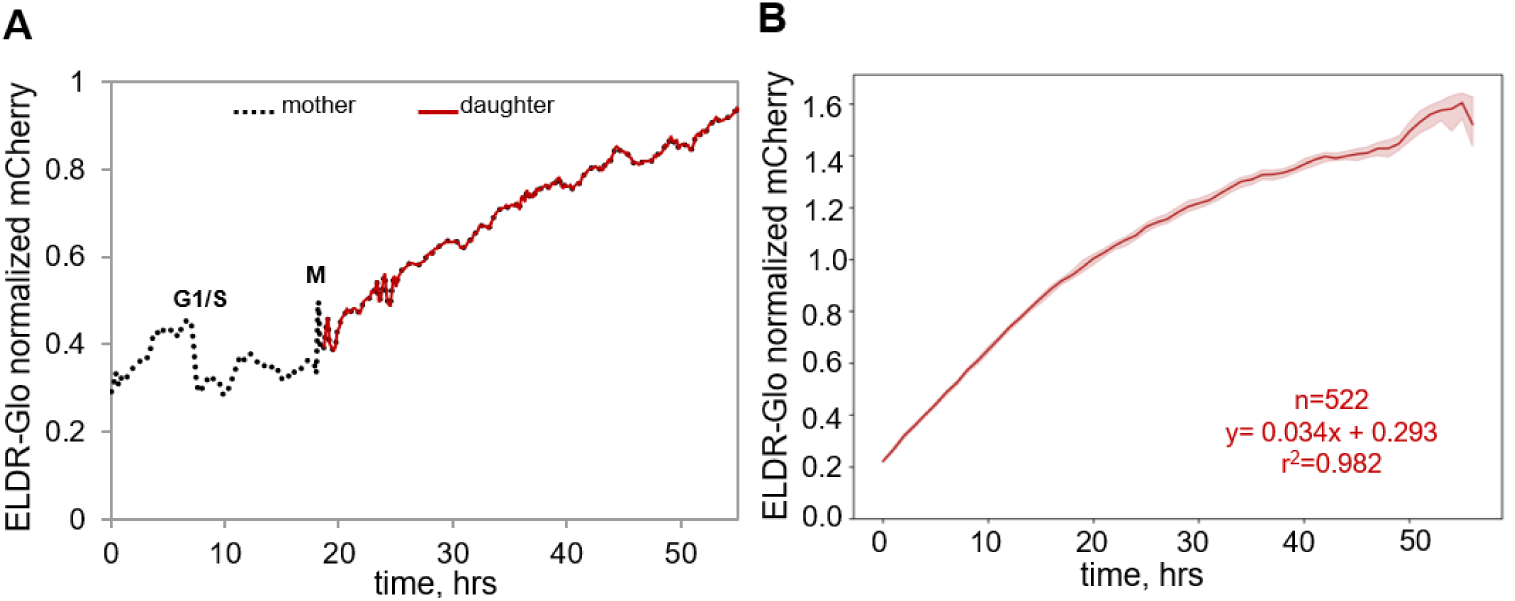
ELDR-Glo dynamics by live cell imaging of arresting cells. (A) Single-cell mother–daughter fluorescence trace showing normalized mCherry intensity (mCherry/mTurquoise) for an RPE cell expressing ELDR-Glo treated with 1 µM Palbociclib. (B) Median normalized mCherry accumulation traces for ELDR-Glo in 522 tracked arresting cells as in (A). Shaded regions represent the 95% confidence interval.

### Compatibility with multiplex immunofluorescence imaging

We next sought to combine the ELDR-Glo biosensor with additional molecular analyses of proliferation and arrest markers. We tested if the mCherry fluorescence can be inactivated prior to antibody staining to allow re-use of the red fluorescence channel. Treating ELDR-Glo expressing cells with 6% hydrogen peroxide efficiently eliminated mCherry fluorescence while preserving nuclear morphology and overall cellular integrity (Figure 4A and Supplementary Figure S5). Of note, the mTurquoise2 signal persisted following peroxide bleaching, indicating that the treatment selectively removed the mCherry biosensor. We and others have previously observed that the copper(I) and ascorbic acid click chemistry reaction often used to detect EdU inactivates both mVenus and GFP because of oxidative damage (*21, 36*). We considered that this effect could be an advantage to inactivate multiple biosensors in fixed cells for sequential analyses. As a test, we imaged fixed cells expressing two cyclin dependent kinase activity biosensors encoding red (CDK4 (*25*)) or yellow-green (CDK2 (*24*)) fluorescent fusions, then treated first with the copper-ascorbic acid solution, washed, and treated with peroxide. We successfully inactivated both the red and green fluorescence and reduced, but did not eliminate blue fluorescence from mTurquoise2 (Supplementary Figure S5). A description of these inactivation steps is provided in Methods.

**Figure 4.**
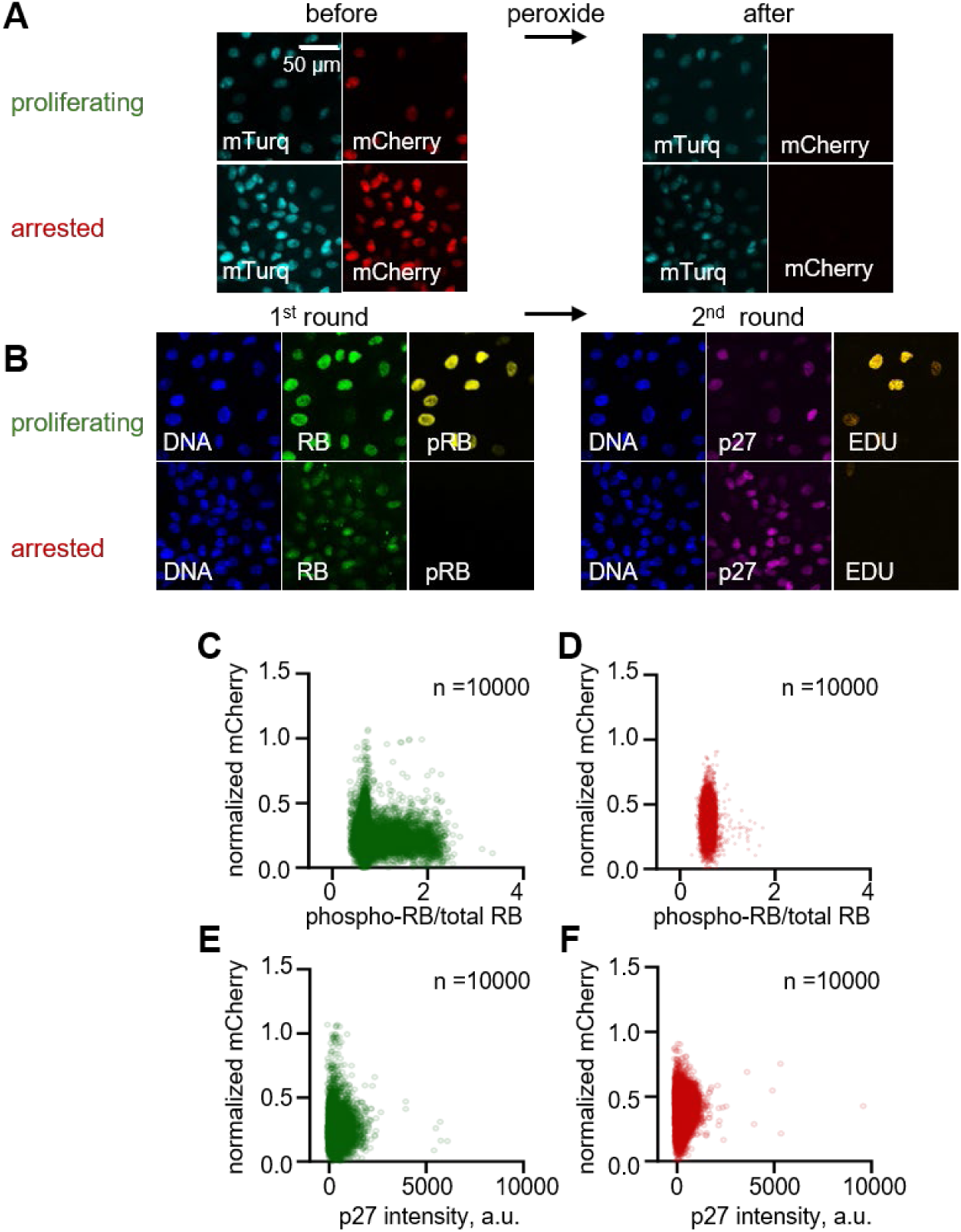
Compatibility with multiplex immunofluorescence imaging. (A) ELDR-Glo fluorescence in proliferating or arrested RPE cells (contact inhibited, 48 hours) before and after bleaching for 15 minurwa with 6% hydrogen peroxide. (B) The same cells in A stained for endogenous RB protein and phospho-RB (S807/S811). Antibodies were eluted (see Methods), then cells stained again for endogenous p27 followed by EdU detection. (C, D) Plot of ELDR-Glo normalized mCherry *vs*. phospho-RB/total RB in 10,000 randomly selected proliferating (C) or arrested (D) cells from A and B detected by indirect immunofluorescence. (E, F) After antibody elution, ELDR-Glo normalized mCherry *vs.* endogenous p27 in the same cells from A-D.

We next combined ELDR-Glo imaging, chemical inactivation, and iterative immunofluorescence for endogenous proliferation biomarkers. We fixed proliferating cells or cells that were arrested from 48 hours of contact inhibition. We collected images of ELDR-Glo signal, then treated cells to bleach the mCherry signal (Figure 4A, before and after). We then stained cells with antibodies to total retinoblastoma protein (RB) and phosphorylated RB (pRB), and we stained for DNA content. After imaging the same field of cells again, we removed the primary and secondary antibodies with strong elution buffer (*37, 38*). We visually confirmed antibody removal and then stained the cells in a second round of immunofluorescence for endogenous p27, a CDK inhibitor protein, and we detected EdU by click chemistry. As expected in proliferating conditions, most, but not all, cells were positive for phosphorylated RB. Proliferating cells had both high and low p27, and the low p27 cells were also low for ELDR-Glo and EdU-positive since both p27 and the biosensor are degraded in S phase (*39*) (Figure 4B, top row). In contrast, arrested cells had more uniformly high ELDR-Glo signal, more uniform p27 levels, and low phospho-RB (Figure 4B, bottom row). Interestingly, in arrested RPE cells p27 abundance did not increase above the maximum G1 level, suggesting that high p27 is not a biomarker to distinguish G0 from G1 in RPE cells. Thus ELDR-Glo can be integrated with immunofluorescence and other detection workflows for relative cell age estimation and molecular analysis in the same cells.

### Compatibility with flow cytometry

We also tested ELDR-Glo performance in both analytical flow cytometry and cell sorting experiments. We first analyzed the mCherry fluorescence in proliferating RPE cells co-stained for DNA content. Fluorescence was highest in G1 cells, near background in S phase cells, and slightly increased in G2 cells, consistent with the live-cell imaging (Figure 5A). Previous reports established that a fraction of cells spontaneously exit the cell cycle and dephosphorylate RB phosphorylation even under proliferation-inducing conditions (*24, 37, 40*). To identify these cells, we fixed and co-stained ELDR-Glo cells for phosphorylated RB and observed that the brightest ELDR-Glo cells were also phospho-RB negative (top ∼15% mCherry signal; Figure 5B). Furthermore, when we arrested cells by contact inhibition (RPE) or CDK4/6 inhibition (MCF-7 cells), ELDR-Glo signal substantially increased (Figure 5C and Supplementary Figure S6).

**Figure 5.**
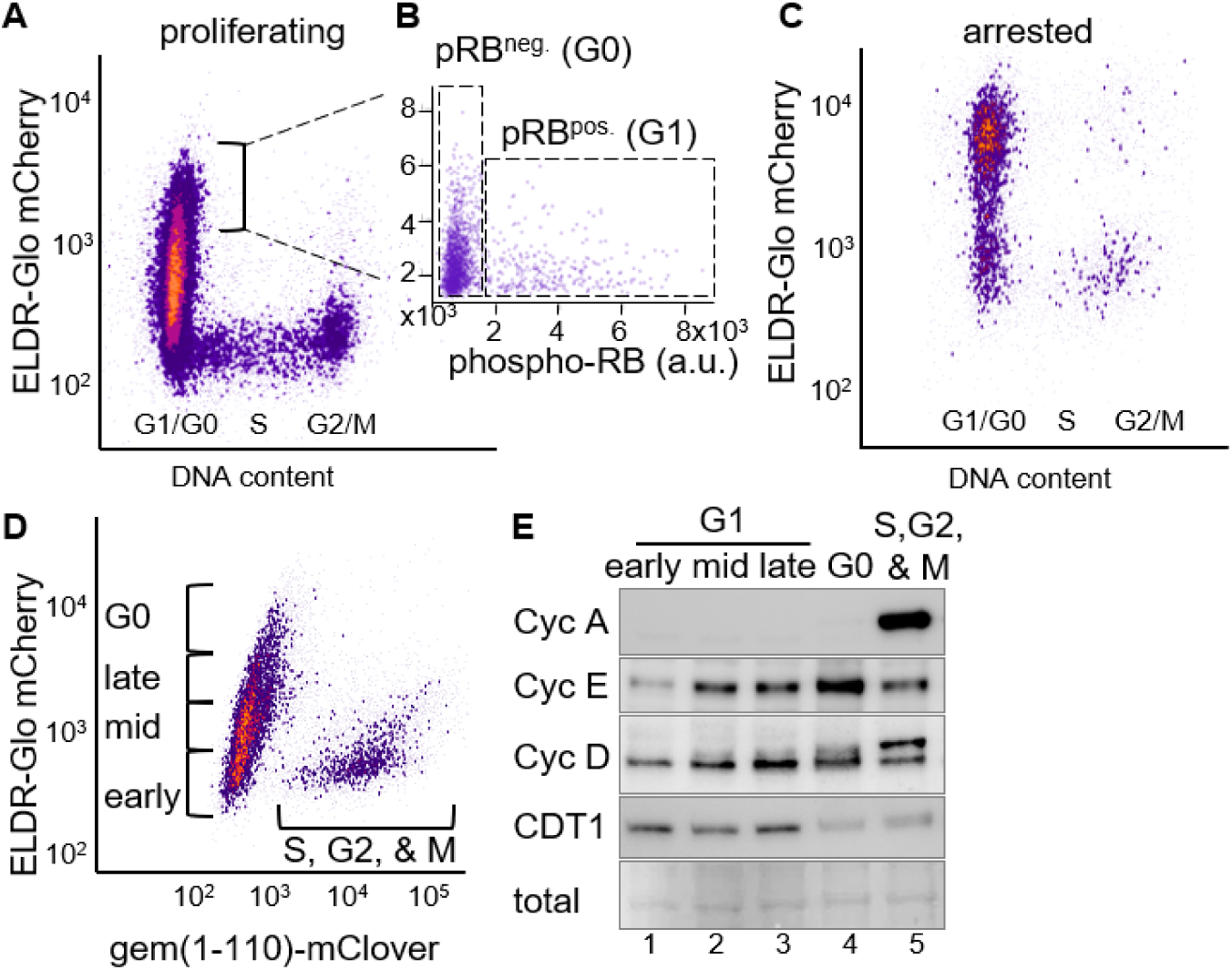
ELDR-Glo compatibility with fluorescence-activated cell sorting (FACS). (A) ELDR-Glo fluorescence detection by analytical flow cytometry in proliferating, fixed RPE cells, co-stained for DNA content with DAPI. (B) The same cells in A were co-stained with phospho-RB (S807/S811). Antibody intensity in the top 15% brightest mCherry cells marked as pRB^neg^ (G0) or pRB^pos^ (G1). (C) ELD R-Glo fluorescence detection by analytical flow cytometry in contact-inhibited RPE cells, co-stained for DNA content with DAPI. (D) Scheme for sorting live mCherry^pos^/mClover^neg^ (early, mid, late G1, and G0) or mCherry^neg^/mClover^pos^ (S, G2, & M) cells. (E) Immunoblotting for the indicated endogenous cell cycle markers in the sorted subpopulations from (D). Ponceau S staining for total protein content.

The wide range of mCherry signal in the G1 population facilitated physically sorting live, unperturbed, asynchronously proliferating cells into early, middle, and late G1 cells plus the G0 subpopulation. To distinguish G1/G0 cells from the other cell cycle phases, we co-expressed a fluorescent fusion of geminin (amino acids 1-110) that is similar to the original cell cycle “FUCCI“ system; geminin is degraded in G1/G0 but increases from S phase through M phase (Figure 5D) (*22, 41*) We subjected these sorted subpopulations to immunoblot analysis for several markers of cell cycle progression (Figure 5E). The various protein abundances matched the expected patterns for cycling untransformed human epithelial cells (*42, 43*): Cyclin A was absent from G1 and G0 cells (lanes 1-4) and abundant in S, G2, and M (lane 5), cyclin D1 and cyclin E1 increased from early to late G1 (lanes 1-3), and CDT1 was down-regulated in S phase (lane 5).

### ELDR-Glo as an indicator of relative quiescence depth

Quiescence is a graded state in which young G0 cells are more likely to re-enter the cell cycle than older G0 cells (*16*). Cells that have been in quiescence longer are thus likely to be more deeply quiescent. Because ELDR-Glo signal correlates with time in G0, we tested its signal relative quiescence depth. A commonly-used biomarker for cell cycle arrest is the loss of endogenous Ki67 expression, a nuclear protein that is present only in proliferating cells. Ki67 is gradually lost in the early stages of quiescence, and Ki67 intensity has been suggested as a proxy for early quiescence depth (*44*). For comparison, we analyzed endogenous Ki67 by immunostaining in proliferating and arrested ELDR-Glo cells. In proliferating cells, both Ki67 and ELDR-Glo signals varied with cell cycle phase as expected (Figure 6A). In arrested cells however, Ki67 levels were mostly very low, near the threshold of detection. In contrast, ELDR-Glo signal continued to accumulate during arrest, producing a substantially broader distribution of normalized mCherry intensities in the arrested population (Figure 6B and Supplementary Figure S7). Thus ELDR-Glo provides a measure of G0 that is complementary to Ki67 but with greater dynamic range for distinguishing differences among quiescent cells of different ages.

**Figure 6.**
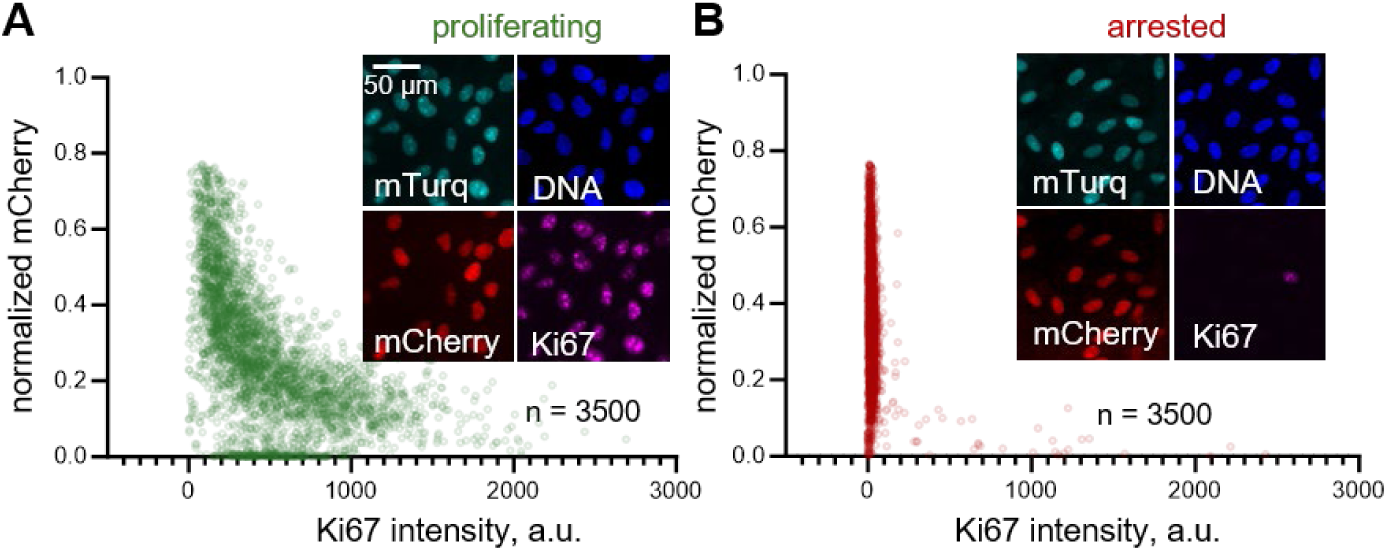
ELDR-Glo signal relative to Ki67 staining. (A) Ki67 immunofluorescence intensity relative to ELDR-Glo signal in proliferating RPE cells. (B) As in A, but RPE cells were arrested with 1 µM palbociclib for 48 hours. X-axis ranges were set below 0 for visualization. Insets are representative fields of cells.

Quiescence depth is defined by how rapidly and/or robustly cells return to proliferation. In general, cells that have been quiescent longer are more deeply quiescent than cells that had more recently exited the cell cycle, suggesting that cell age correlates with quiescence depth. To test this idea, we analyzed ELDR-Glo intensity in assays that reveal differences in quiescence depth. We arrested cells by contact inhibition, then released them by replating at low cell density. We immediately began imaging them, then tracked ELDR-Glo dynamics for the next 40 hours. We determined when each cell entered S phase from the rapid drop in mCherry intensity (Figure 7A). A subset of cells remained arrested for the full time of imaging with modest increases in ELDR-Glo signal over time. Other cells also increased biosensor signal at a similar rate but then entered S phase at different times after release. We analyzed the ELDR-Glo signal at the start of imaging relative to the time of S phase entry for each individual tracked cell. This analysis revealed a general and consistent correlation between lower ELDR-Glo starting signal and earlier S phase (Supplementary Figure S8).

**Figure 7.**
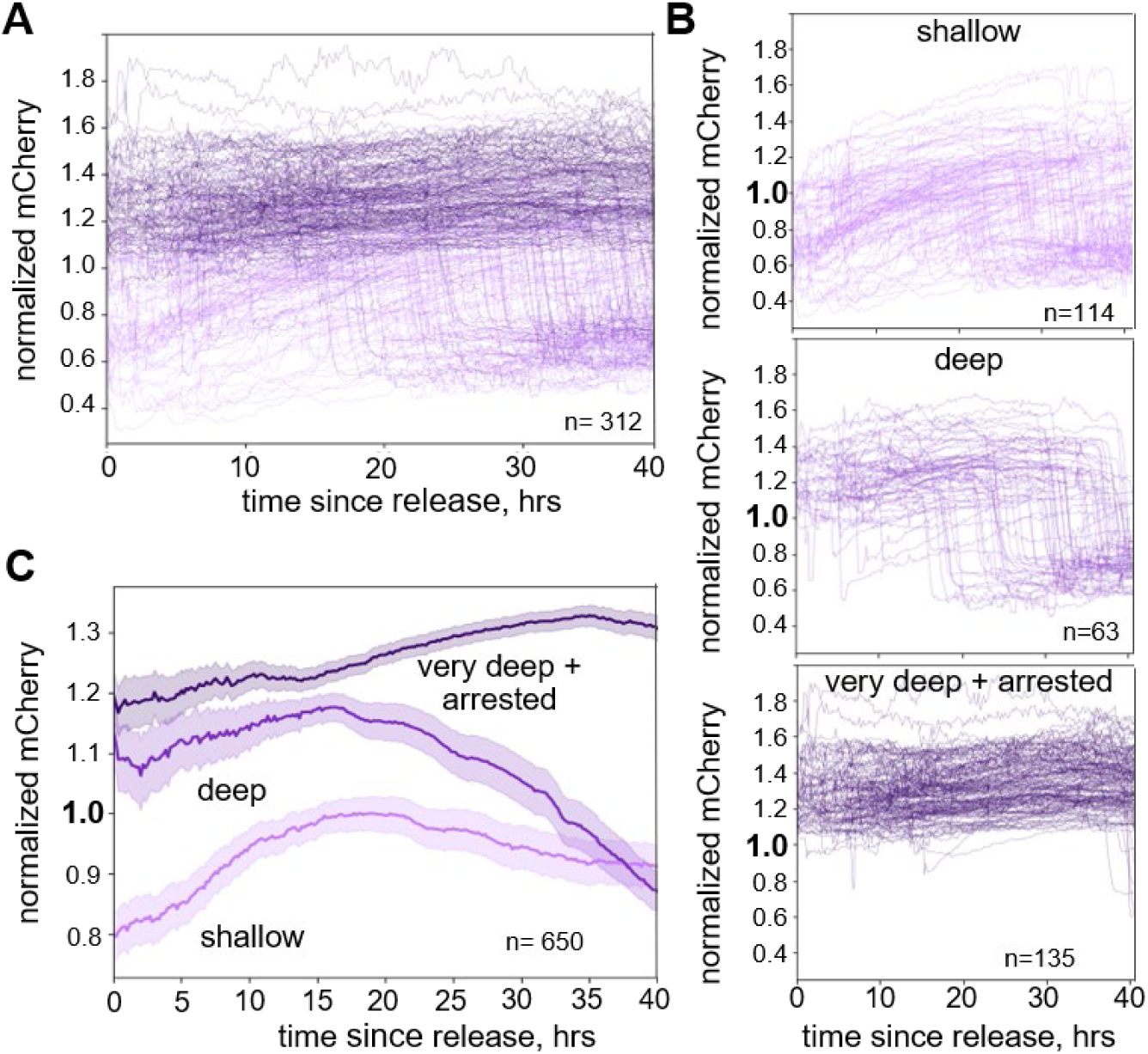
ELDR-Glo as a relative quiescence depth indicator. (A) ELDR-Glo RPE cells were arrested by contact inhibition for 48 hours then replated at low density, followed by immediate live cell imaging and continuous tracking. A random subset of the total 650 traces is shown. (B) Cells were classified by a starting ELDR-Glo signal below 1.0 as shallow or above 1.0 as deep. Cells that did not enter S within 40 hours were classified as very deep + arrested. Individual traces of a subset of the three cell classifications are plotted. (C) Complete tracks of the three groups of cells plotted as mean + 95% confidence interval.

We divided cells into three groups: cells that entered S phase with ELDR-Glo signal at the time of release from G0 above 1.0 (presumed deep), cells that started below 1.0 (presumed shallow), and cells that did not enter S phase within the 40 hours of imaging. Cells in this last group might have entered S phase after 40 hours or remained arrested, so we marked them as “very deep + arrested”. For visualization, we separately plotted ELDR-Glo signal over time for each of the three groups (Figure 7B). Strikingly, most of the cells with ELDR-Glo signal below 1.0 at the time of release entered S phase earlier than those that started above 1.0. Shallow quiescent cells began S phase an average of 13.4 ± 8.0 hours after release, whereas deep quiescent cells began S phase 26.2 ± 10.3 hours after release. The differences in S phase entry for the groups classified by mean ELDR-Glo signal at the time of release from quiescence suggest their designations as shallow or deep was appropriate.

We also plotted the mean ELDR-Glo signal over time for the shallow, deep, and very deep/arrested cells (Figure 7C). For all three groups, the mean biosensor signal initially increased with time as cell age continued to increase, but then the mean dropped as more and more cells in the group entered S phase. The mean signal dropped earlier for the shallow quiescent cells than for the more deeply quiescent cells (Figure 7C and replicate in Supplementary Figure S9). By definition, the mean signal drop for very deep/arrested would have begun after 40 hours. These differences are also consistent with the low ELDR-Glo cells starting from a shallow quiescent state relative to the high ELDR-Glo cells in the same population. Importantly, ELDR-Glo signal at the time of re-stimulation correlated with *future* quiescence depth phenotype. In other words, different ELDR-Glo signals among the quiescent cells while they were still quiescent generally matched their actual quiescence depth. This feature contrasts with other quiescence depth experiments that require re-stimulation to infer how deeply quiescent cells had been before stimulation.

Because biosensor signal increases with time since the last S phase, and ELDR-Glo signal indicates relative age among cells in both G1 and G0, we determined how well the biosensor can predict cell age. To test this idea, we imaged live cells that were arresting from CDK4/6 inhibition, and we randomly selected frames from the video. Using linear regression for the relationship between signal and cell age (Figure 3B), we used ELDR-Glo to classify 500 cells from these frames as either older or younger than 10 hours. We then compared these age predictions to their true cell age from the tracked live cell data. By this test, the biosensor correctly predicted more than 86% of the cells as either older or younger than 10 hours (Figure 8A).

**Figure 8.**
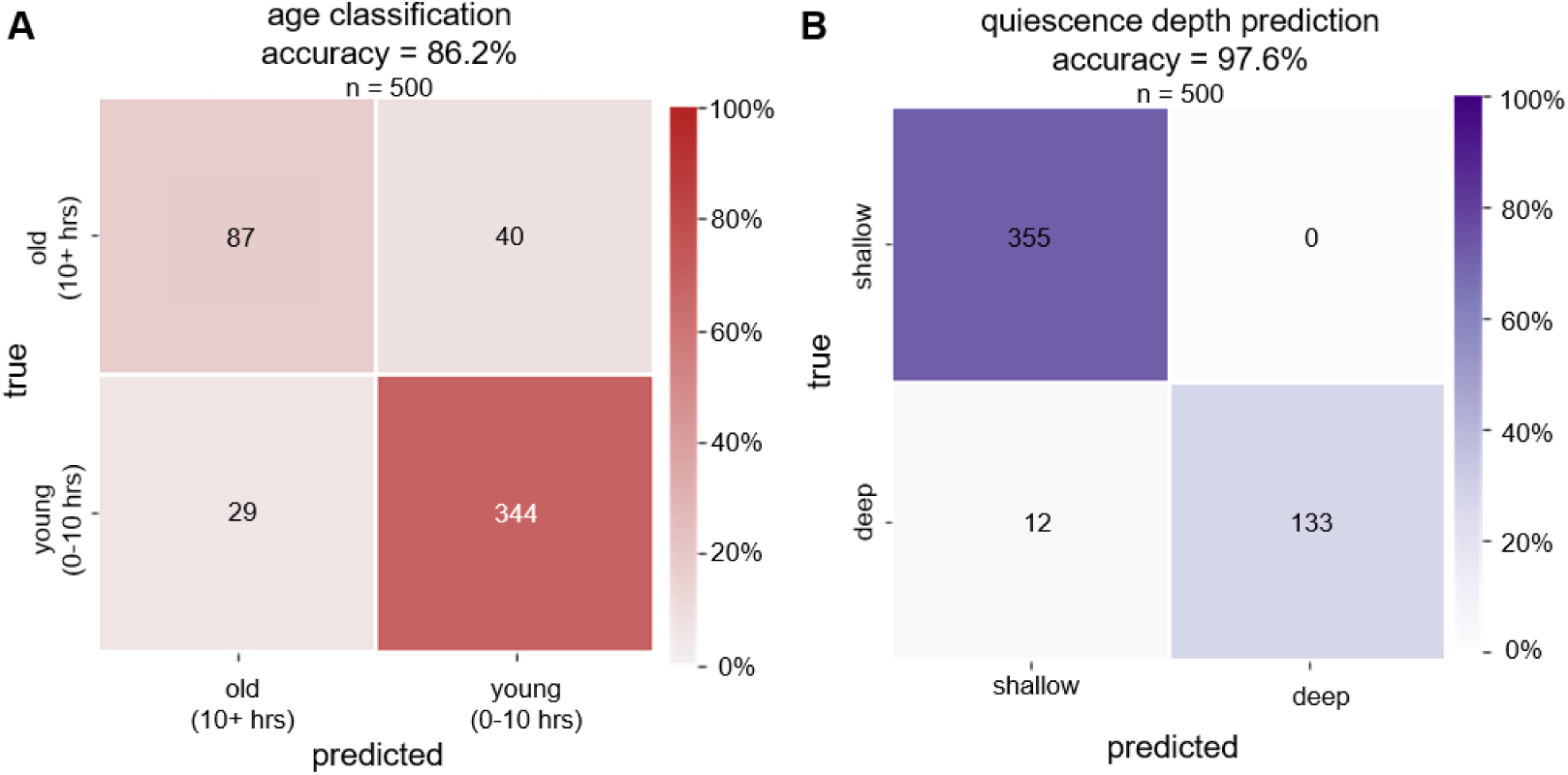
Predicting relative age and quiescence depth from ELDR-Glo signals. (A) Randomly-selected drug-induced RPE cells in different frames of a live cell movie were classified as older or younger than 10 hours by ELDR-Glo signal alone. The true age of each cell was determined, and the relative accuracy of the age classification was calculated (86.2%). The table is color-coded by the accuracy of the predictions. (B) Cells were contact inhibited for 48 hours, then released by replating at low density and subjected to live cell imaging for 40 hours (same cells as Figure 7). Immediately after they had attached and spread, cells were classified as either shallow or deep by ELDR-Glo signal alone as in Figure 7B. The time of S-phase entry was determined from the live cell imaging, and the relative accuracy of the predictions was calculated (97.6%). The table is color-coded by the accuracy of the predictions.

Finally, we tested if ELDR-Glo signal accurately predicts deep or shallow quiescence behavior. We assigned 500 contact-inhibited cells from an independent replicate as likely to be in deep or shallow quiescence using biosensor signal at the time of release above or below 1.0, respectively. We compared these predictions to the true behavior after release by live cell imaging. The biosensor accurately predicted relative quiescence depth for more than 97% of the cells (Figure 8B). Together, these tests indicate that the ELDR-Glo biosensor is a quantitative measure of relative cell age which correlates quiescence depth and can likely predict the responsiveness of quiescent cells to proliferative cues.

## Discussion

A challenge for studies of proliferation and cell cycle arrest dynamics has been the need for long-term imaging or, alternatively, artificial synchronization and samples collected at different times. Synchronization introduces perturbations such as stress responses that could influence cell behavior and associated molecular markers (*45–48*). The ELDR-Glo biosensor is applicable to large numbers of cells of different relative ages under otherwise unperturbed conditions. ELDR-Glo should also be informative in settings where live cell imaging is not feasible, such as *in vivo* analyses of most tissues or tumors.

Chemical bleaching of the fluorescent timer (or other related reporters) allows iterative antibody labeling to integrate age measurements with molecular markers in the same cells. ELDR-Glo is presumably also compatible with fluorescence in situ hybridization (FISH) or other staining methods. In addition, ELDR-Glo can be used with flow cytometry to physically isolate cells based on their relative age. ELDR-Glo could readily be used for downstream molecular profiling, including transcriptomic, metabolomic, and proteomic analyses.

Standard proliferation biomarkers such as DNA synthesis or phospho-RB staining distinguish proliferation from arrest but do not distinguish among different arrested cells. Although Ki67 is gradually downregulated after the final cell division, this signal falls below the threshold of detection relatively quickly (*44*). As a result, biologically distinct shallow and deep quiescent cells at later points are often indistinguishable in fixed samples despite differences in functional potential. The ELDR-Glo biosensor addresses this issue because its signal increases rather than decreases with time in arrest. An alternative G0 reporter based on p27 accumulation in arrested cells was described previously, suggesting that p27 levels might be a general indicator of cell age and thus, quiescence depth (*49*). However, neither the reporter nor endogenous p27 accumulates much higher than a typical G1 maximum in either RPE cells or primary vascular endothelial cells (HUVEC). Moreover, p27 increases in contact-inhibited HUVEC but actually decreases during quiescence induced by laminar flow (*50*), meaning p27 is not a universal marker of G0. Others have also reported substantial differences in molecular markers during different kinds of arrest (*51, 52*). In contrast, ELDR-Glo dynamics are based on an intrinsic property of the mCherry variant, making it generally agnostic to cell type and method of arrest. The slow dynamics of mCherry maturation also gives ELDR-Glo a relatively large dynamic range.

Because ELDR-Glo signal correlates with age, and thus quiescence depth, it serves as a probable indicator of future proliferative competence in the arrested cells while they are still arrested. We consider this to be an important advantage for quiescence depth studies since users can identify which cells are likely more deeply or shallowly quiescent *in situ*. Assessing depth usually requires re-stimulating cells and then determining how deeply quiescent cells had been before stimulation, but at that point, they are no longer quiescent. Alternative methods would require live imaging during entry into arrest to directly measure how old cells are or comparing average measurements in populations arrested for different amounts of time. Such bulk analyses have been instrumental in establishing quiescence depth as an important biological phenomenon, but ELDR-Glo provides a unique approach to analyzing the quiescent cells themselves.

Several considerations are important when interpreting ELDR-Glo signals. First, the biosensor reports time since the last S phase rather than time after mitosis or quiescent cell state *per se*. In the cell types we analyzed, G2 phase is typically very short, so nearly all signal accumulation is after division in G1 or G0. However, any prolonged arrest in G1 would allow biosensor signal to increase in a manner similar to deeply quiescent cells (with the possible exception of very high DNA damage). Moreover, in systems with long G2 phases, ELDR-Glo accumulation becomes a measure of time in G2. In that regard because the CDT1 PIP degron is sensitive to levels of DNA damage well beyond the natural replication stress in unperturbed S phase (*33, 34*), the biosensor might report dynamics following DNA repair, although this idea would require testing.

Second, the dynamic range is constrained by the kinetics of fluorescent protein maturation and eventual signal plateau. As the rate of signal increase slows, resolution among the older cells decreases. At the highest biosensor signals, we can only conclude that all of the bright cells are older than the age of cells at the time of signal plateau. If this property is relevant for a given study, we recommend pilot calibrations to determine when the signal plateaus for the imaging setup in use.

Third, although normalization with the mTurquoise component reduces variability, especially in polyclonal populations, the relationship between biosensor signal and absolute cell age is imperfect. ELDR-Glo is a relative measure of cell age rather than an exact clock. When we subdivided RPE cells older than 10 hours into additional categories in Figure 8A, the prediction accuracy dropped. If determining actual age is critical, calibration curves with live imaging or time courses within the cells being tested is important. Nonetheless, within a certain range, ELDR-Glo is very accurate at predicting which cells are older and more deeply quiescent than neighboring young and shallow quiescent cells.

ELDR-Glo can link cell age to proliferation potential. Additional possible applications include analyzing tissue stem cell quiescence, investigating tumor cell dormancy and therapy resistance, and identifying factors that modulate quiescence depth in regeneration and disease contexts. In addition, the ability to stratify cells by age without live imaging opens new opportunities for high-throughput screening and multiomic profiling of defined cell populations. The ELDR-Glo biosensor adds a new tool for studying aspects of intercellular heterogeneity that have previously been difficult to measure.

## Materials and Methods

### Plasmid vectors

The entire ELDR-Glo construct was synthesized by Twist Bioscence and inserted into pLenti CMV Blast DEST (706–1) (*53*). The insert is in the following order from 5’-3’ with short linkers between elements (in italics): human CDT1 amino acids 1-17 (MEQRRVTDFFARRRPGP-*RTRG*), SV40 nuclear localization sequence (PKKKRKV-*GAQ*) double HA tag (YPYDVPDYAGYPYDVPDYA-*TS*), mCherry “medium” variant from Subach et al. 2009 (*31*), T2A ribosome skipping sequence (*LE*-GSGEGRGSLLTCGDVEENPGP-*GS*), double SV40 nuclear localization sequence (PKKKRKV-*AQ*-PKKKRKV-*LVPVAT*), mTurquoise2. Plasmid sequences were verified by restriction digest and sequencing; the ELDR-Glo plasmid will be deposited at Addgene. The geminin 1-110-mClover vector was acquired from Addgene (# 83915) (*41*); PCNA-mTurquoise has been described (*21*) and is available at Addgene (#118617).

### Cell culture and stable cell lines

HEK293T, RPE1-hTERT, and MCF7 cell lines were obtained from ATCC and confirmed to be mycoplasma negative. HEK293T and RPE1-hTERT cells were maintained in Dulbecco’s modified Eagle medium (Sigma-Aldrich Chemistry) supplemented with 10% fetal bovine serum (FBS) (Avantor Seradigm), 2 mM L-glutamine (Gibco), and 100 U/mL penicillin–streptomycin (Gibco). MCF7 cells were cultured in Eagle’s Minimum Essential Medium (Gibco) supplemented with 10% FBS and 10 μg/mL human recombinant insulin (Gibco). All cells were maintained at 37°C in a humidified incubator with 5% CO₂ and passaged using 0.25% trypsin-EDTA. Human umbilical vein endothelial cells (HUVEC) were cultured according to the manufacturer’s recommendations in EBM2 (Lonza; No. CC-3162) with growth factors (Lonza; No. CC-3162, EGM-2) at 37 °C, 5% CO2 and used between passages 2-6.

To generate RPE and MCF7s cell lines: lentiviral expression plasmids were co-transfected with ΔNRF and VSVG (gifts from J. Bear) virus packaging plasmids into HEK293T cells using 50 μg/mL polyethylenimine (PEI)-Max (Sigma-Aldrich Chemistry). Two days later, virus-containing supernatant was collected and used to transduce cells in the presence of 8 μg/mL polybrene (Millipore) for at least 6 hours (maximum 24 hours). In some cases, a second round of viral transduction was performed. Cells that successfully integrated the transduced constructs were selected by treating with blasticidin (Sigma-Aldrich Chemistry).

To generate ELDR-Glo-expressing primary HUVEC: for lentivirus production, HEK293T cells were seeded at 4.5 x 10^6^ cells/10 cm dish in antibiotic-free DMEM supplemented with 10% FBS for 24h, then co-transfected with the ELDR-Glo plasmid, the psPAX2 packaging plasmid, and the pMD2.G envelope plasmid (Addgene plasmid # 12260 and #12259, gifts from D. Trono) using the Calcium Phosphate Transfection Kit (Takara Bio, #631312). After overnight incubation, the medium was replaced with HUVEC medium.

Viral supernatants were collected 48–72 h post-transfection, clarified by centrifugation and passed through a 0.45 µm Polyethersulfone (PES) filter, aliquoted, and stored at -80 °C until use. Early passage (P2-P4) HUVEC were plated at 3x10^5^ cells/well in a 6-well plate in 1 mL medium/well. 500 µL of lentiviral supernatant was added directly to each well 30 minutes after seeding, and medium was replaced after 24 h. After 24 h of additional recovery, cells were selected with 10 µg/mL of blasticidin S (ThermoFisher) and maintained in selective medium for 2 passages.

All cell lines generated are polyclonal except for the RPE1-hTERT expressing ELDR-Glo. For clonal isolation, cells were plated sparsely following viral transduction, and clones were screened for expression of the construct of interest. Cells were finally enriched by fluorescence-activated cell sorting (BD FACSymphony S6 Cell Sorter (Biosciences)) of live cells using both mTurquoise and mCherry intensities.

### Cell treatments

For DNA synthesis detection, cells were treated with 10 µM EdU for 30 minutes prior to harvest (flow cytometry) or fixation (4i). EdU was detected as described below. For quiescence depth analysis, RPE cells were seeded at high density and allowed to grow until there were very few mitotic cells in the dish followed by additional incubation for 48 hours. To release from quiescence, cells were trypsinized and re-plated at low density into glass-bottom well plates (Cellvis) and immediately subjected to live-cell imaging for up to 50 hours. For drug arrests, cells were treated with the indicated concentrations of palbociclib (PD0332991 CDK4/6i, Selleckchem). DNA damage cells were treated with Neocarzinostatin (NCS,Sigma) at varying concentrations of 0-200 nM for 4 hours, followed by a wash with 1x PBS.

### Antibodies

Primary antibodies and their working dilutions: retinoblastoma protein RB, Cell Signaling Technology Cat# 9309 (Ms, 1:500 for immunofluorescence); phospho-retinoblastoma protein pRB, Cell Signaling Cat#8516 (1:200 for flow cytometry; 1:1000 for immunofluorescence); p27, Cell Signaling Technology Cat #3686 (1:800 for immunofluorescence); Ki67, Abcam Cat. #15580 (1:1000 for immunofluorescence); 53BP1, Santa Cruz Cat# sc-22760 (1:1000 for immunofluorescence); anti-HA tag, Santa Cruz Biotechnology, Cat# sc-515312 (1:1000 for immunoblotting); mouse anti-cyclin A2, Cell Signaling Technology, Cat# 4656 (1:2000 for immunoblotting); mouse anti-cyclin E1, Cell Signaling Technology, Cat# 4129 (1:2000 for immunoblotting); mouse anti-cyclin D1, Santa Cruz Biotechnology Cat# sc-8396, (1:1000 for immunoblotting), and rabbit anti-CDT1, Cell Signaling Technology, Cat# 8064 (1:1000 for immunoblotting).

Secondary antibodies: Donkey anti-mouse AF555, ThermoFisher Cat# A32790 (1:500 for immunofluorescence), AF488 Azide, ThermoFisher AF10266 (1:1000 for EdU detection); HRP-conjugated donkey anti-rabbit IgG, Jackson ImmunoResearch, Cat# 711-035-152 (1:10,000 for immunoblotting), and HRP-conjugated donkey anti-mouse IgG, Jackson ImmunoResearch Cat# 715-035-150 (1:10,000 for immunoblotting).

### Immunoblotting

Cells were collected by trypsinization and lysed on ice for 20 minutes in CSK buffer (100 mM NaCl, 300 mM sucrose, 3 mM MgCl₂, and 10 mM PIPES, pH 7.0) supplemented with 0.5% Triton X-100, protease inhibitors (1 µg/mL pepstatin A, 0.1 mM AEBSF, 1 µg/mL aprotinin, and 1 µg/mL leupeptin), and phosphatase inhibitors (1 mM β-glycerophosphate, 10 µg/mL phosvitin, and 1 mM sodium orthovanadate). Lysates were clarified by centrifugation and quantified using a Bradford assay (Bio-Rad) with bovine serum albumin as a standard. Proteins (typically 20 µg per lane) in sample buffer (1% SDS, 2.5% 2-mercaptoethanol, 0.1% bromophenol blue, 50 mM Tris pH 6.8, 10% glycerol) were boiled for 5 minutes, centrifuged, and resolved on SDS–polyacrylamide gels then transferred to polyvinylidene difluoride membranes (ThermoFisher). Membranes were stained with Ponceau S, imaged for total protein, then rinsed and blocked for one hour in 5% milk in Tris-buffered saline with 0.1% Tween-20 (TBST) and incubated with primary antibodies overnight at 4°C in 2.5% milk/TBST. Membranes were washed three times for 5 minutes each with TBST, then probed with horseradish peroxidase-coupled secondary antibodies and washed 3 times with TBST. Signals were detected using ECL Prime (Amersham) and collected using a ChemiDoc imaging system (Bio-Rad).

### Flow cytometry

Analytical flow cytometry was performed essentially as described but without pre-extraction with detergent (*54*). Data were collected with an Attune NxT Flow cytometer (ThermoFisher). For ELDR-Glo mCherry fluorescence detection, control samples included untransduced parental RPE1 cells or unstained cells to determine thresholds for positive signals. FCS data files were analyzed in FCS Express 7 software (De Novo Software). Raw single cell values were exported for analysis from FCS Express to GraphPad Prism.

For live cell sorting, single-color control cells were plated and used for gating. Cells were harvested by trypsinization and filtered through 50 μm Celltrics filters (Sysmex), counted, washed once with warm 1X PBS, then resuspended in FluoroBriteTM DMEM (Invitrogen) supplemented with 10% FBS, 2 mM L-glutamine and Pen/Strep. DNA was stained with Vybrant Dye Cycle Violet (Invitrogen) at 1:1000 for ∼1 hour at 37°C. Cells were kept at 37°C until immediately before sorting. Cell sorting was performed as five-way sorting on the BD FACSymphony S6 Cell Sorter (Biosciences) at room temperature. Sorted cells were pelleted, washed with 1X PBS, then flash frozen and stored at -80°C for later use.

### Live-cell imaging

Before imaging RPE and MCF7s, cells were seeded at low density in glass-bottom plates (Cellvis Cat#P96-1.5P) in appropriate imaging media (no phenol red; RPE: FluoroBrite DMEM (Invitrogen Cat#A1896701), MCF7s: supplemented with FBS, L-glutamine, and penicillin/streptomycin). For contact inhibition, cells were seeded at a high density.

HUVEC (P6) were seeded at 2x10^5^ cells/well (high density) in a 24-well fibronectin-coated glass bottom plate in EBM2 (Lonza, Cat# CC-3162) with added growth factors (Bullet kit, CC-3162, Lonza, referred to as EGM2), and incubated for 2 hours prior to imaging.

All live-cell fluorescence imaging was performed using a Nikon Ti Eclipse inverted microscope equipped with Plan Apochromat objectives (20×, numerical aperture 0.75) and a Hamamatsu Orca Flash 4.0s CMOS camera. Environmental conditions were maintained using an Okolabs incubation chamber at 37°C under 5% CO₂ with controlled humidity. Focus was maintained using the Nikon Perfect Focus System throughout the duration of imaging. Fluorescence images were acquired using standard filter sets for CFP, YFP, and DsRED. The following filter sets were used (excitation; beam splitter; emission filter; Chroma): CFP (425-445/455/465-495nm), YFP (490-510/515/520-550nm), and DsRED (540-580/585/593-668). Image acquisition was performed using NIS-Elements AR software at 10-minute intervals, with 2x2 stitching. Exposure times and illumination intensities were optimized to minimize photobleaching and phototoxicity, and no detectable effects on cell behavior were observed under these conditions. Stage coordinates were recorded to enable longitudinal tracking of individual cells and alignment with downstream analyses.

### Chemical bleaching of fluorophores

RPE cells expressing PCNA-mTurquoise2, DHB fragment-mVenus (*24*), and RB fragment-mCherry (*25*) were seeded at low density in glass-bottom plates, incubated for ∼24 hours, then fixed using 4% PFA, washed, and permeabilized. mCherry signals were quenched using 6% hydrogen peroxide diluted in distilled-deionized water (ddH_2_0) for 15-30 minutes. mVenus was inactivated in fixed and washed cells by treatment with 1 mM CuSO₄ (Sigma) and 100 mM ascorbic acid (Fisher) in ddH_2_O for 15-30 minutes. After either bleaching treatment, samples were extensively washed 3x with 1x PBS.

### Iterative Indirect Immunofluorescence Imaging (4i) and fixed cell microscopy

Iterative antibody staining was performed essentially as described in Stallaert *et al.* 2022 (*37*). Fixed cells were incubated in 4i blocking solution (100 mM maleimide (Sigma), 100 mM NH4Cl (Sigma) and 1% bovine serum albumin in 1x PBS) for 1 hour and incubated with primary antibodies diluted as indicated in conventional blocking solution (1% bovine serum albumin in 1x PBS) overnight at 4°C. Samples were rinsed 3x with PBS and then incubated in secondary antibodies and 1:1000 DAPI for 1 hour with gentle shaking, then rinsed 3x with PBS and imaged in imaging buffer (700 mM N-acetyl-cysteine (Sigma) in ddH2O, pH 7.4). After imaging, cells were rinsed 3x with ddH2O and incubated with elution buffer (0.5 M L-glycine (Sigma), 3 M urea (Sigma), 3 M guanidine chloride (ThermoFisher Scientific), and 70 mM TCEP-HCl (Sigma) in ddH20 adjusted to pH 2.5) 3x for 10 minutes with shaking. Cells were then blocked and incubated with the second round of primary antibodies. EdU incorporation was detected after the final immunostaining by reaction with 488 azide (1:1000) in PBS with 1 mM CuSO4 (Sigma) and 100 mM ascorbic acid (ThermoFisher) fresh in ddH2O for 30 minutes at room temperature in the dark.

Fluorescence imaging was performed using a Nikon Ti Eclipse inverted microscope equipped with Plan Apochromat objectives (20×, numerical aperture 0.75) and a Hamamatsu Orca Flash 4.0s CMOS camera. The following filter sets were used (excitation; beam splitter; emission filter; Chroma): DAPI (383–408/425/435-485 nm), CFP (425-445/455/465-495nm), YFP (490-510/515/520-550nm), DsRED (540-580/585/593-668) and Cy5 (590-650/660/663-738nm). Image acquisition was performed using NIS-Elements AR software with 6x6 stitching. Micrographs were saved as TIFF files and imported into ImageJ. Where necessary for clear visualization, any adjustments to brightness and contrast were applied equally to the entire image.

### Image processing, segmentation, and tracking

Live cell tracking and iterative indirect immunofluorescence analysis were accomplished using modified Python scripts from Zikry et *al.* (*55*). Time-lapse image sequences and immunostained images were corrected for background and shading artifacts. Nuclear regions of interest were segmented based on fluorescence intensity using automated thresholding and watershed-based separation. In cases where automated segmentation was insufficient, nuclei were manually outlined. Fluorescence intensities for mCherry and mTurquoise were quantified within the same nuclear regions of interest. The ELDR-Glo signal was calculated as the ratio of mCherry to mTurquoise intensity to normalize for expression differences across cells. Where applicable, entry into S phase was defined by the appearance of characteristic punctate nuclear PCNA-mVenus foci (*26*).

### Statistical analysis

Statistical analyses were performed using GraphPad Prism and Python scripts. Comparisons between groups were conducted using ANOVA with appropriate post hoc tests. Live-cell imaging experiments included three technical replicates and a minimum of two independent biological replicates. FACS and 4i experiments included at least two biological replicates.

Normalized ELDR-Glo signal was plotted as a function of time for individual cells, and linear regression analysis was used to estimate accumulation rates. To evaluate the accuracy of ELDR-Glo-based classification, normalized mCherry intensity was converted to predicted cellular age using a linear calibration model derived under drug-induced conditions and live cell imaging (normalized mCherry = m × age + b). Predicted age was obtained by inverting this relationship and was binarized into early (≤ 10 hour) and late (>10 hour). For quiescence depth classification, a threshold for normalized mCherry intensity was defined based on distribution-based binning in an independent replicate. This threshold for starting ELDR-Glo signal (normalized mCherry = 1.08 arbitrary units) was then applied to an independent dataset to classify cells as shallow or deeply quiescent.

Confusion matrices were generated by comparing predicted and observed classifications. Classification accuracy was calculated as the fraction of correctly classified cells across all categories. All analyses were performed using custom Python scripts, which are available to the public.

## Data visualization

Data visualization was performed using Python (v3.13) with matplotlib (v3.11.1) and seaborn (v0.13.2), as well as GraphPad Prism (v11).

## Acknowledgments

The authors are grateful to the members of the Cook lab for advice and helpful discussions, to Sam Wolff, Arwa Sattar, Arthi Adavikolanu, and Shruti Bommareddy for technical support, and to Jeffrey Jones for managerial assistance. Clover-Geminin(1–110) was a gift from Michael Lin (Addgene plasmid # 83915). pLenti CMV Blast DEST (706–1) was a gift from Eric Campeau & Paul Kaufman (Addgene plasmid # 17451). Lentiviral packaging plasmids were a gift from J. Bear.

## Funding

NIH National Institute of General Medical Sciences (NIGMS) grant R35GM141833 (JGC)

NIH National Cancer Institute (NCI) grant R61CA302625 (JGC)

NIH National Institute of Diabetes and Digestive Kidney Diseases grant T32DK007737 (MSJ)

American Heart Association (AHA) 23PRE1027147 (DF)

NIH National Heart, Lung, and Blood Institute grant R35HL139950 (VLB) NIH P30 CA016086 (Lineberger Comprehensive Cancer Center support grant)

The UNC Flow Cytometry Core Facility (RRID:SCR_019170) and the UNC Proteomics Core Facility are supported in part by P30 CA016086 Cancer Center Core Support Grant to the UNC Lineberger Comprehensive Cancer Center. Research reported in this publication was supported in part by the North Carolina Biotech Center Institutional Support Grant 2017-IDG and by the NIH 1UM2AI30836-01.

## Author contributions

Conceptualization: MSJ VLB JGC

Methodology: MSJ JGC DF LM

Investigation: MSJ SK DF TH LM ND ML WA

Visualization: MSJ SK DF LM

Supervision: VLB JGC

Writing—original draft: JGC MSJ SK WA DF

Writing—review & editing: MSJ VLB JGC

### Data and materials availability

All data are available in the main text or the supplementary materials or from the author upon reasonable request.

### Supplementary Materials

Figures S1-S9

## References

1. K. Vermeulen, D. R. Van Bockstaele, Z. N. Berneman, The cell cycle: a review of regulation, deregulation and therapeutic targets in cancer. Cell Proliferation 36, 131–149 (2003).

2. J. P. Matson, J. G. Cook, Cell cycle proliferation decisions: the impact of single cell analyses. FEBS J 284, 362–375 (2017).

3. D. Hanahan, R. A. Weinberg, Hallmarks of cancer: the next generation. Cell 144, 646–674 (2011).

4. H. K. Matthews, C. Bertoli, R. A. M. De Bruin, Cell cycle control in cancer. Nature Reviews Molecular Cell Biology 23, 74–88 (2022).

5. R. J. Duronio, Y. Xiong, Signaling pathways that control cell proliferation. Cold Spring Harb Perspect Biol 5, a008904 (2013).

6. C. T. J. van Velthoven, T. A. Rando, Stem Cell Quiescence: Dynamism, Restraint, and Cellular Idling. Cell stem cell 24, 213–225 (2019).

7. E. A. Robinson, A. R. Barr, G0 or no-G0: phosphatase control of quiescence and cell cycle entry. Biochem Soc Trans 54, 585–599 (2026).

8. O. Marescal, I. M. Cheeseman, Cellular Mechanisms and Regulation of Quiescence. Dev Cell 55, 259–271 (2020).

9. M. Mitra, S. L. Batista, H. A. Coller, Transcription factor networks in cellular quiescence. Nat Cell Biol 27, 14–27 (2025).

10. M. S. Johnson, J. G. Cook, Cell cycle exits and U-turns: Quiescence as multiple reversible forms of arrest. Fac Rev 12, 5 (2023).

11. A. Eames, S. Chandrasekaran, Leveraging metabolic modeling and machine learning to uncover modulators of quiescence depth. PNAS Nexus 3, pgae013 (2024).

12. B. Liu et al., Extracellular Fluid Flow Induces Shallow Quiescence Through Physical and Biochemical Cues. Front Cell Dev Biol 10, 792719 (2022).

13. A. de Morree, T. A. Rando, Regulation of adult stem cell quiescence and its functions in the maintenance of tissue integrity. Nat Rev Mol Cell Biol 24, 334–354 (2023).

14. K. Fujimaki et al., Graded regulation of cellular quiescence depth between proliferation and senescence by a lysosomal dimmer switch. Proc Natl Acad Sci U S A 116, 22624–22634 (2019).

15. A. G. Evertts et al., H4K20 methylation regulates quiescence and chromatin compaction. Mol Biol Cell 24, 3025–3037 (2013).

16. L. H. Augenlicht, R. Baserga, Changes in the G0 state of WI-38 fibroblasts at different times after confluence. Exp Cell Res 89, 255–262 (1974).

17. J. S. Kwon et al., Controlling Depth of Cellular Quiescence by an Rb-E2F Network Switch. Cell Rep 20, 3223–3235 (2017).

18. S. He, N. E. Sharpless, Senescence in Health and Disease. Cell 169, 1000–1011 (2017).

19. J. K. Freitas Leal, M. J. W. Adjobo-Hermans, R. Brock, G. Bosman, Acetylcholinesterase provides new insights into red blood cell ageing in vivo and in vitro. Blood Transfus 15, 232–238 (2017).

20. K. T. Soh, J. D. Tario, Jr., K. A. Muirhead, P. K. Wallace, Probing cell proliferation: Considerations for dye selection. Methods Cell Biol 186, 1–24 (2024).

21. G. D. Grant, K. M. Kedziora, J. C. Limas, J. G. Cook, J. E. Purvis, Accurate delineation of cell cycle phase transitions in living cells with PIP-FUCCI. Cell Cycle 17, 2496–2516 (2018).

22. A. Sakaue-Sawano et al., Visualizing spatiotemporal dynamics of multicellular cell-cycle progression. Cell 132, 487–498 (2008).

23. A. Sakaue-Sawano et al., Genetically Encoded Tools for Optical Dissection of the Mammalian Cell Cycle. Mol Cell 68, 626–640 e625 (2017).

24. S. L. Spencer et al., The proliferation-quiescence decision is controlled by a bifurcation in CDK2 activity at mitotic exit. Cell 155, 369–383 (2013).

25. H. W. Yang et al., Stress-mediated exit to quiescence restricted by increasing persistence in CDK4/6 activation. eLife 9, (2020).

26. H. Leonhardt et al., Dynamics of DNA replication factories in living cells. J Cell Biol 149, 271–280 (2000).

27. S. Xie, S. Zhang, G. De Medeiros, P. Liberali, J. M. Skotheim, The G1/S transition in mammalian stem cells in vivo is autonomously regulated by cell size. Nature Communications 16, (2025).

28. R. C. Adikes et al., Visualizing the metazoan proliferation-quiescence decision in vivo. eLife 9, (2020).

29. S. Xie, J. M. Skotheim, A G1 Sizer Coordinates Growth and Division in the Mouse Epidermis. Curr Biol 30, 916–924 e912 (2020).

30. S. Pauklin, L. Vallier, The cell-cycle state of stem cells determines cell fate propensity. Cell 155, 135–147 (2013).

31. F. V. Subach et al., Monomeric fluorescent timers that change color from blue to red report on cellular trafficking. Nat Chem Biol 5, 118–126 (2009).

32. C. G. Havens, J. C. Walter, Docking of a specialized PIP Box onto chromatin-bound PCNA creates a degron for the ubiquitin ligase CRL4Cdt2. Mol Cell 35, 93–104 (2009).

33. T. Senga et al., PCNA is a cofactor for Cdt1 degradation by CUL4/DDB1-mediated N-terminal ubiquitination. J Biol Chem 281, 6246–6252 (2006).

34. J. Hu, Y. Xiong, An evolutionarily conserved function of proliferating cell nuclear antigen for Cdt1 degradation by the Cul4-Ddb1 ubiquitin ligase in response to DNA damage. J Biol Chem 281, 3753–3756 (2006).

35. S. F. Pereira, R. L. Gonzalez, Jr., J. Dworkin, Protein synthesis during cellular quiescence is inhibited by phosphorylation of a translational elongation factor. Proc Natl Acad Sci U S A 112, E3274–3281 (2015).

36. A. Löschberger, T. Niehörster, M. Sauer, Click chemistry for the conservation of cellular structures and fluorescent proteins: ClickOx. Biotechnology Journal 9, 693–697 (2014).

37. W. Stallaert et al., The structure of the human cell cycle. Cell Syst 13, 230–240 e233 (2022).

38. G. Gut, M. D. Herrmann, L. Pelkmans, Multiplexed protein maps link subcellular organization to cellular states. Science 361, (2018).

39. A. C. Carrano, E. Eytan, A. Hershko, M. Pagano, SKP2 is required for ubiquitin-mediated degradation of the CDK inhibitor p27. Nat Cell Biol 1, 193–199 (1999).

40. A. J. Pulianmackal et al., Monitoring Spontaneous Quiescence and Asynchronous Proliferation-Quiescence Decisions in Prostate Cancer Cells. Front Cell Dev Biol 9, 728663 (2021).

41. B. T. Bajar et al., Fluorescent indicators for simultaneous reporting of all four cell cycle phases. Nature methods 13, 993–996 (2016).

42. C. Rega et al., High resolution profiling of cell cycle-dependent protein and phosphorylation abundance changes in non-transformed cells. Nat Commun 16, 2579 (2025).

43. S. Gookin et al., A map of protein dynamics during cell-cycle progression and cell-cycle exit. PLoS Biol 15, e2003268 (2017).

44. I. Miller et al., Ki67 is a Graded Rather than a Binary Marker of Proliferation versus Quiescence. Cell Rep 24, 1105–1112 e1105 (2018).

45. J. P. Matson et al., Intrinsic checkpoint deficiency during cell cycle re-entry from quiescence. J Cell Biol 218, 2169–2184 (2019).

46. H. A. Coller, What’s taking so long? S-phase entry from quiescence versus proliferation. Nat Rev Mol Cell Biol 8, 667–670 (2007).

47. S. Cooper, K. Shedden, Microarray analysis of gene expression during the cell cycle. Cell Chromosome 2, 1 (2003).

48. M. J. Beyrouthy et al., Identification of G1-regulated genes in normally cycling human cells. PLoS One 3, e3943 (2008).

49. T. Oki et al., A novel cell-cycle-indicator, mVenus-p27K-, identifies quiescent cells and visualizes G0-G1 transition. Sci Rep 4, 4012 (2014).

50. N. T. Tanke et al., Endothelial Cell Flow-Mediated Quiescence Is Temporally Regulated and Utilizes the Cell Cycle Inhibitor p27. Arterioscler Thromb Vasc Biol 44, 1265–1282 (2024).

51. H. A. Coller, L. Sang, J. M. Roberts, A new description of cellular quiescence. PLoS Biol 4, e83 (2006).

52. M. Min, S. L. Spencer, Spontaneously slow-cycling subpopulations of human cells originate from activation of stress-response pathways. PLoS Biol 17, e3000178 (2019).

53. E. Campeau et al., A versatile viral system for expression and depletion of proteins in mammalian cells. PLoS One 4, e6529 (2009).

54. D. L. Bolhuis et al., USP37 prevents unscheduled replisome unloading through MCM complex deubiquitination. Nat Commun 16, 4575 (2025).

55. T. M. Zikry et al., Cell cycle plasticity underlies fractional resistance to palbociclib in ER+/HER2− breast tumor cells. Proceedings of the National Academy of Sciences 121, e2309261121 (2024).

