## Supplementary Figures S1-S9 for "ELDR-Glo, a biosensor for cell age that correlates with quiescence depth"

Martha S. Johnson, *et. al.*

**This PDF file includes:**

Figs. S1 to S9

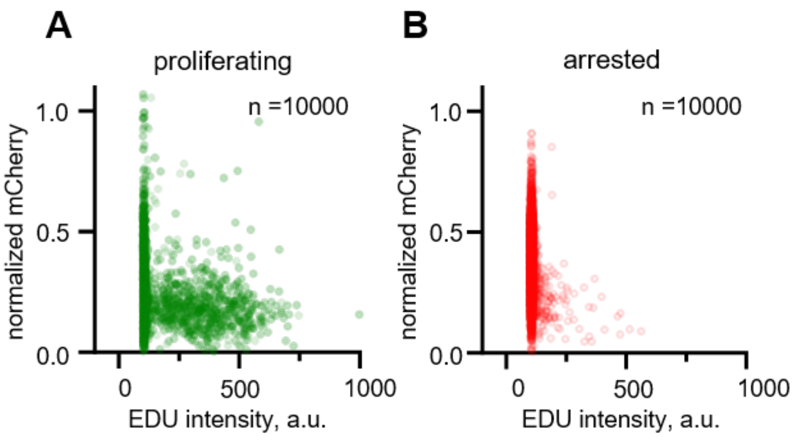


**Fig. S1. ELDR-Glo in fixed cells**

(A) ELDR-Glo signal and DNA synthesis to identify S-phase cells by EdU incorporation in proliferating RPE cells

(B) As in A but in RPE cells arrested by contact inhibition for 48 hours.

Both datasets were randomly down-sampled to 10,000 cells for visualization.

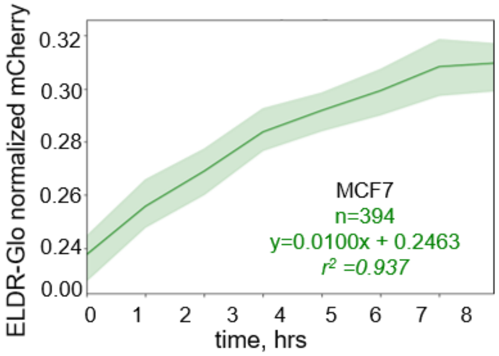


**Fig. S2.ELDR-Glo G1 dynamics in proliferating MCF7s cells.**

Median normalized mCherry signal in G1 phase of 394 tracked MCF7 cells. Tracks were aligned to the frame after cell division (0 hrs). Shaded regions represent the 95% confidence interval.

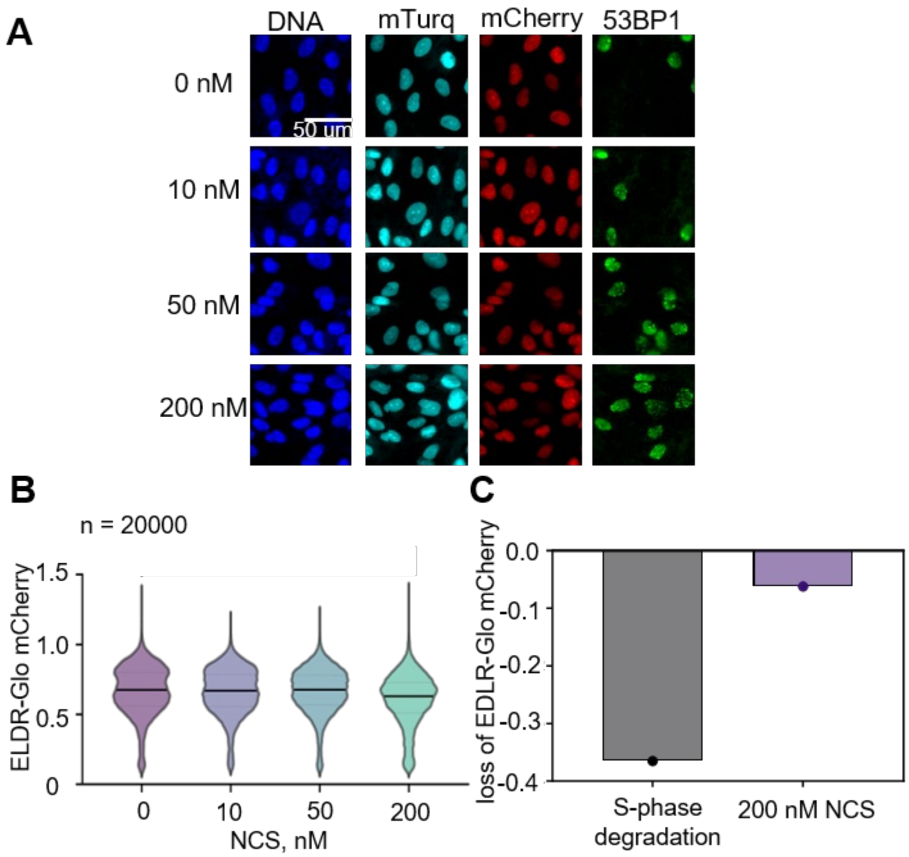


**Fig. S3. ELDR-Glo is resistant to moderate DNA damage.**

RPE cells were treated with 0, 50, 100, and 200 nM NCS for 4 hrs to induce double-strand DNA breaks.

(A) Cells were fixed and stained for DNA, imaged for the mTurquoise2, ELDR-Glo mCherry and endogenous 53BP1 localization by immunostaining.

(B) Normalized mCherry signal in NCS-treated cells after the indicated doses; 20,000 cells per condition.

(C) Fold-change in mean mCherry signal in S phase cells or after 200 nM NCS treatment.

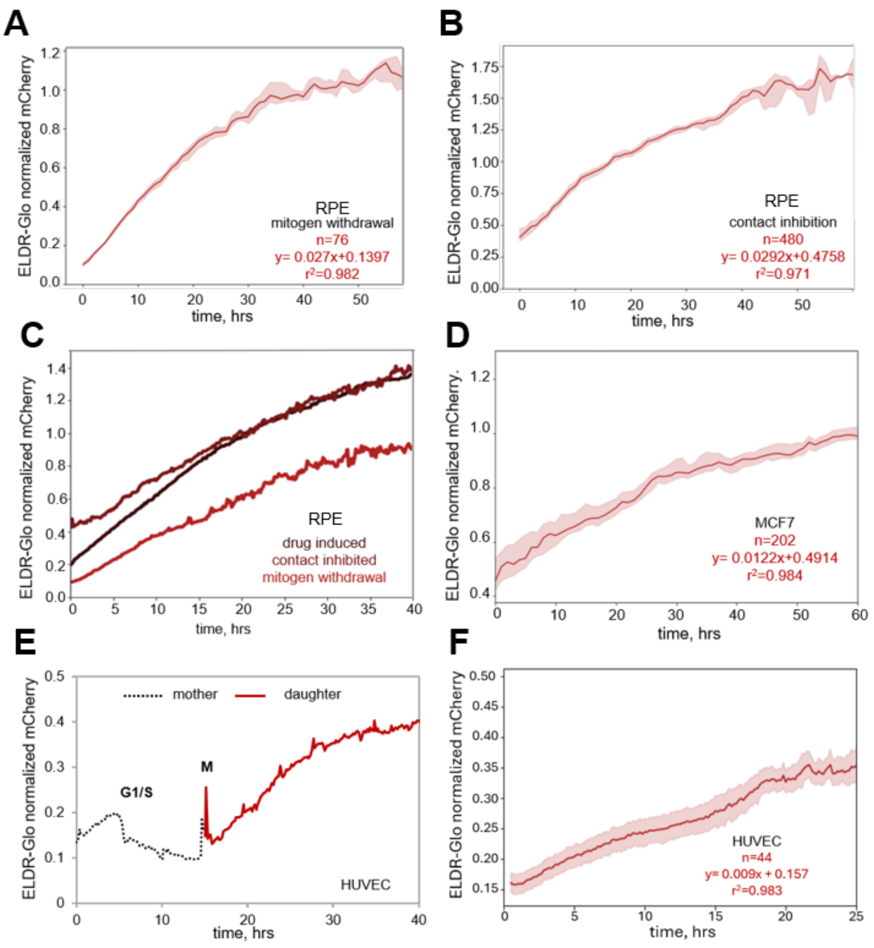


**Fig. S4. ELDR-Glo dynamics by live cell imaging of arresting cells.**

(A) Median normalized mCherry accumulation for ELDR-Glo in RPE cells during mitogen starvation. Cells were tracked from cell division. Shaded regions represent the 95% confidence interval. n=76 cells

(B) As in A except that RPE cells were tracked during arrest from contact inhibition. n=480 cells

(C) Median normalized mCherry accumulation of arrest from CDK4/6 inhibition (data from main Figure 3B), contact inhibition, and mitogen withdrawal.

(D) Median normalized mCherry accumulation for ELDR-Glo in MCF7s cells arrested with 1 μM palbociclib. Cells were tracked from cell division. Shaded regions represent the 95% confidence interval. n=202 cells

(E) Primary HUVEC expressing ELDR-Glo were imaged as they approached quiescence from contact inhibition. One mother–daughter fluorescence trace showing normalized mCherry intensity (mCherry/mTurquoise).

(F) Median normalized mCherry accumulation traces for ELDR-Glo in 45 tracked HUVEC arresting from low mitogen treatment. Shaded regions represent the 95% confidence interval. n=44 cells

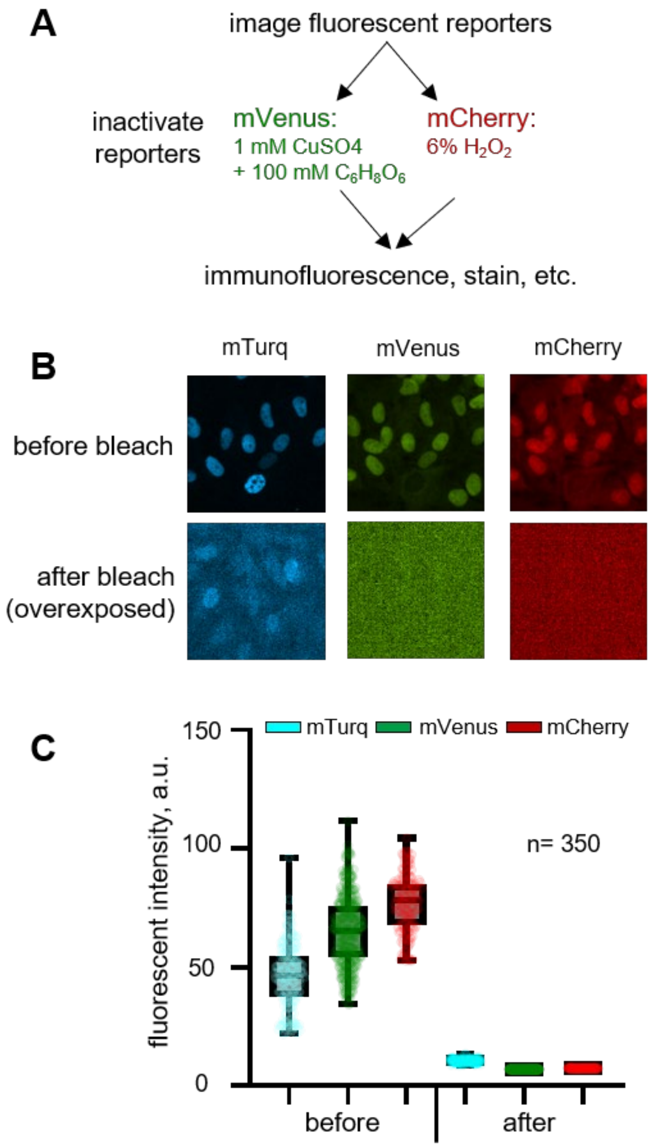


**Fig S5: Bleaching fluorescent biosensors for subsequent immunostaining.**

(A) Workflow for chemical bleaching of mCherry and mVenus (or GFP). See methods for details. For experiments in which both mCherry and mVenus were inactivated in the same cells, cells were treated with CuS0₄ + ascorbic acid for 30 minutes to inactivate mVenus, washed with 1x PBS, and then treated with 6% hydrogen peroxide for 15 or 30 minutes to inactivate the mCherry.

(B) Cells expressing PCNA-mTurquoise, CDK1/2 activity mCherry sensor (Spencer, *et al.*, 2013), and CDK4/6-mVenus activity (Yang, *et al.,* 2020) were fixed, imaged for the reporters, and then treated to inactivate the reporters. Representative images before and after treatment are shown; post-bleach images were overexposed.

(C) Quantification of reporter fluorescence intensities before and after bleaching; n= 350 cells.

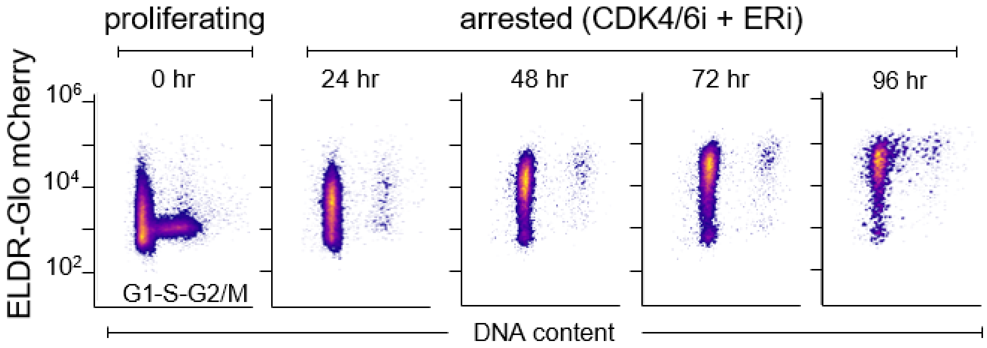


**Fig. S6. ELDR-Glo accumulation in arrested MCF-7 cells.**

The MCF-7 breast cancer cell line was transduced with the ELDR-Glo construct, and one clone selected. Cells were treated with a combination of 200 nM palbociclib (CDK4/6 inhibitor) and 20 nM fulvestrant (estrogen receptor antagonist) for the indicated times then analyzed by flow cytometry for ELDR-Glo mCherry intensity and DNA content (DAPI).

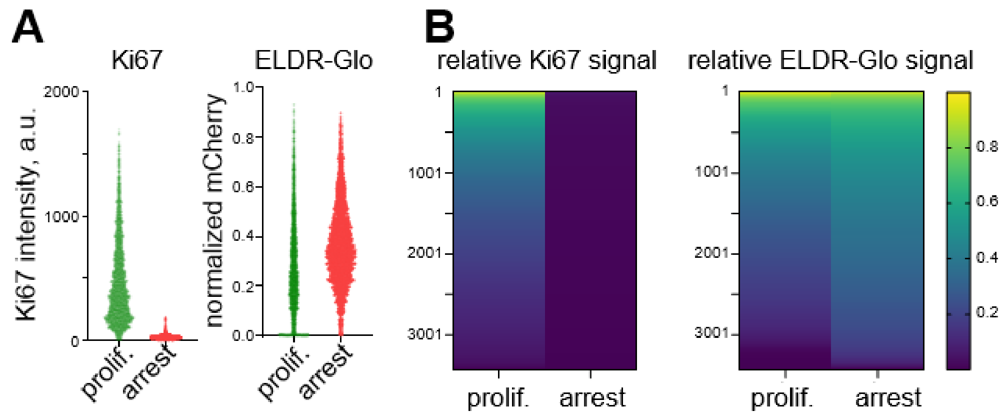


**Fig. S7. Dynamic range of ELDR-Glo in arrested cells.**

(A) Ki67 immunostaining intensities and ELDR-Glo signals from Figure 6 are plotted for proliferating and arrested cells.

(B) Signal intensities for each cell were calculated relative to the maximum Ki67 or ELDR-Glo signal, respectively, excluding the top and bottom 1%.

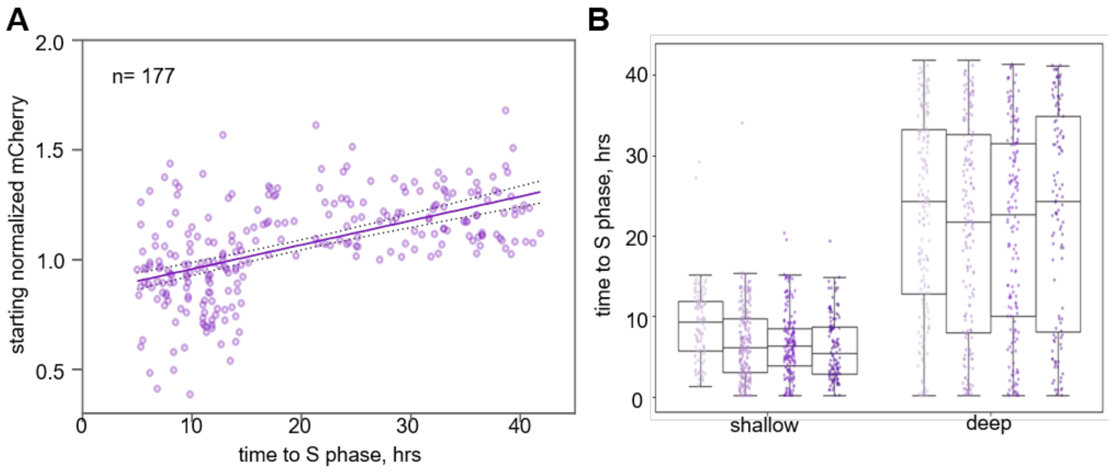


**Fig. S8. ELDR-Glo in quiescence depth experiments.**

(A) Starting signal vs S phase entry time from Figure 7B.
(B) S phase entry for four replicates of the experiment in (A).

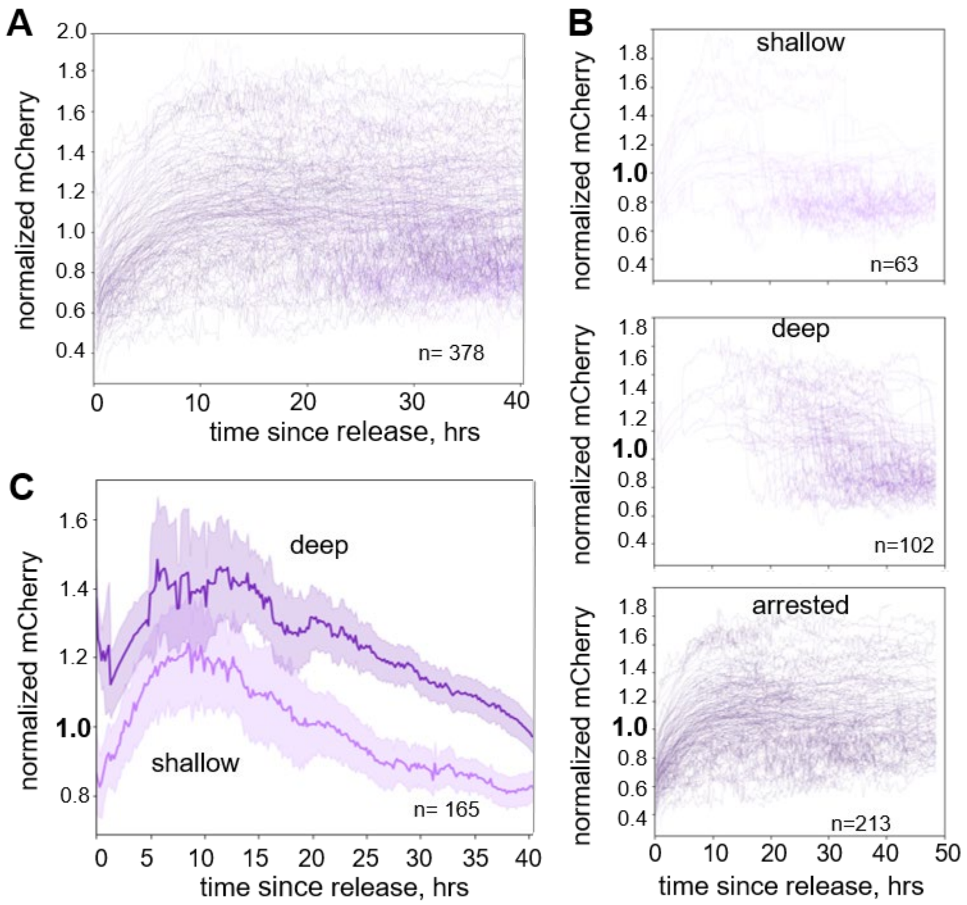


**Fig. S9. Biological replicate quiescence depth experiment.**

(A) ELDR-Glo- bearing RPE cells were arrested by contact inhibition for 72 hrs then replated at low density, followed by immediate live cell imaging and continuous tracking. A random subset of the total 378 traces is shown.

(B) Cells were classified by a starting ELDR-Glo signal below 1.0 as shallow or above 1.0 as deep. Cells with no S phase within 40 hrs were classified as arrested. Individual traces of a subset of the three cell classifications are plotted.

(C) Complete tracks of the shallow and deep cells in B plotted as median + 95% confidence interval.
